# Exercise-induced DNA damage response and memory formation in mice

**DOI:** 10.1101/2025.09.26.678909

**Authors:** Natalie Ricciardi, Robyn McCartan, Juliana Laverde Paz, Huifen Cao, Caroline Borja, Ava Cherry, Alvin Baranov, Clarence Locklear, Olga Wójcikowska, Zane Zeier, Philipp Kapranov, Claes Wahlestedt, Claude-Henry Volmar

## Abstract

This study reveals that acute aerobic exercise enhances memory formation through a controlled DNA damage mechanism, offering crucial insights into Alzheimer’s disease (AD) prevention. This work challenges the traditional view that DNA damage is inherently harmful, demonstrating that minor, reversible DNA single-strand breaks (SSBs) induced by exercise serve as necessary primers for memory consolidation – a mechanism that may be impaired in AD pathogenesis. AD affects 1 in 9 adults over 65, with ~95% being late-onset cases where up to 40% of risk factors are modifiable through lifestyle interventions like exercise. While exercise demonstrably lowers AD risk, underlying mechanisms remain unclear. This study provides a missing mechanistic link by showing how exercise-induced DNA damage repair systems could counteract the DNA damage accumulation and repair dysregulation that are established hallmarks of brain aging and AD. In data presented herein, young mice showed significantly higher SSB rates in active genomic regions compared to aged mice, suggesting the decline of a protective mechanism (i.e., hormesis) with aging – potentially explaining increased AD susceptibility in older adults. The present study also suggests that exercise-induced SSBs are not random cellular damage but precisely targeted events that occur at genes essential for neuroplasticity, synaptic function, and memory formation. These breaks activate PARP1 (Poly ADP ribose polymerase 1), a crucial DNA damage sensor that simultaneously initiates repair processes while facilitating transcriptional programs necessary for memory consolidation. This mechanism may represent how exercise “primes” the brain against the pathological DNA damage accumulation seen in AD. In support of this, in behavioral experiments, a single exercise bout converted sub-threshold learning into robust long-term memory formation. This memory enhancement correlated with upregulation of both neurotrophic genes (BDNF, Fos) and DNA repair enzymes (PARP1, PARP2), demonstrating coordinated damage-repair processes that appear compromised in AD. We identify HPF1 as a critical cofactor enabling PARP1 to perform trans-ADP-ribosylation of histones, linking DNA damage sensing to epigenetic chromatin remodeling required for memory-related gene expression. This pathway represents a novel therapeutic target for AD, as restoring efficient DNA repair mechanisms might slow or prevent memory loss.

## Introduction

Alzheimer’s disease (AD), the most common form of dementia, affects 1 in 9 adults age 65+ in the U.S. and has limited therapeutic options that fail to halt disease progression. Late-onset AD (LOAD) or sporadic AD makes up about 95% of AD cases, and of these cases modifiable risk factors account for up to 40%^1^. These risk factors include numerous markers of metabolic health like blood pressure, cholesterol level, and glucose and insulin tolerance which can all be regulated by an increase in physical activity. Both aerobic exercise and resistance training have been shown to lower the risk ratio for developing age-associated AD. However, this finding has not led to the development of novel, disease modifying therapies which are desperately needed. This is, in part, due to the lack of mechanistic understanding behind how physical activity improves memory and brain function in both healthy and diseased individuals.

The cognitive benefits of exercise in AD prevention may be explained through the lens of hormesis, a term used to describe any process by which a stressful stimulus, when received in a specific low-dose range, produces a favorable phenotypic outcome; however, when that range is exceeded, the benefit is reduced or even becomes harmful^2^. Exercise inherently induces hormetic processes through shifts of molecular physiology by raising energy demands to support muscle activation, increasing production of reactive oxygen species (ROS), and inducing cortisol production, among many other shifts in molecular physiology^3,4^. Acute moderate exercise induces a controlled level of stress on a molecular level that leads to favorable adaptations in organ systems throughout the body, including the brain^5,6^. In this context, stress is registered on a cellular level by a decline in the concentration of bioenergetic metabolites like glucose and free fatty acids, a drop in available molecular oxygen, as well as a drop in the ATP:AMP ratio^7,8^. These are just a few of the biochemical changes that, if not experienced in a healthy context, can lead to cell death rather than beneficial adaptations against future insults^9^. There is a mountain of preclinical and clinical evidence that a single bout of exercise has acute effects on memory and cognition^10–14^. Additionally, a growing body of evidence shows that regular exercise training is both preventative and therapeutic for AD^15,16^. It is hypothesized that these cognitive benefits are due, in part, to an eventual increase in BDNF protein expression in the blood and key memory-related regions of the brain. This has made BDNF the gold standard molecular indicator of exercise-induced neuroplasticity^17–20^.

One such underexplored exercise-induced mechanism is DNA damage and repair: specifically, DNA single-strand breaks (SSBs) and repair. Single-stranded DNA breaks are a common form of genomic damage in neurons that disproportionately impact the transcriptionally active regions of genes essential for neuronal physiology. We and others have previously shown that DNA damage accumulation and dysregulation of DNA damage repair (DDR) enzymes are hallmarks of aging in the brain and periphery and are exacerbated in cases of AD^21–27^. While aerobic exercise is proposed as a panacea for aging-associated diseases, the precise mechanisms behind these health benefits are not fully elucidated^28,29^. DDR is a critical process that is responsible for chromatin maintenance and functional longevity of postmitotic neurons^30,31^.

Unlike other cell types, neurons are unable to divide and replace damaged cells and must maintain genomic stability for an entire lifetime while sustaining the high levels of transcription required for learning, memory, and other neuronal processes. The DNA within a cell incurs thousands of daily lesions as a result of homeostatic processes including metabolic reactive oxygen species (ROS) production, transcription initiation, and even as intermediates of DDR^32,33^. Notably, most of these lesions occur as SSBs which are approximately 1000-fold more common than double-stranded DNA breaks (DSBs)^34,35^.

Adaptive SSB breaks occur as a part of transcription initiation and most frequently occur in highly expressed immediate early genes that support synaptic and cellular plasticity. Because these SSBs occur during active transcription, repair must occur concurrently with gene expression. However, these adaptive breaks carry an inherent risk to genomic integrity when they are unable to be properly repaired. Whereas double strand breaks typically elicit strong repair responses, SSBs in actively transcribed regions elicit less of a response. The ability of SSBs to be repaired is especially crucial to brain tissue as evidenced by the development of rare neurological ataxias when multiple SSB repair enzymes are mutated^36–39^.

Over time, repeated adaptive DNA breaks may burden repair systems, particularly with aging as repair efficiency declines. While DNA damage accumulation has typically been associated with brain aging and neurodegeneration, understanding how SSB distribution changes with age is critical for distinguishing between pathological damage and functionally relevant breaks that may support neuroplasticity and cognitive function. Interestingly, exercise induces DNA damage in peripheral tissues throughout the body including muscle, liver, and blood^40–44^. Despite this, there is little research investigating the relationship between exercise-induced DNA damage and the brain^45–47^. Specifically, it is unknown how DNA damage and the concomitant DDR response relate to neuroprotection or how physical activity primes against the neuronal DNA damage that accumulates with age.

Given that SSBs occur preferentially at transcriptionally active regions and require coordinated repair responses, understanding the molecular machinery that detects and responds to these breaks is critical. Poly ADP ribose polymerase 1 (PARP1) serves as a key sensor of DNA damage and coordinator of repair responses, making it a prime candidate for mediating the beneficial effects of exercise-induced DNA damage. This dual role of PARP1 as both a DNA damage sensor and transcriptional coordinator positions it as an ideal mediator of hormetic responses to exercise. When exercise induces controlled SSB formation, PARP1 activation could simultaneously initiate repair processes while facilitating the transcriptional programs necessary for memory consolidation and neuroplasticity.

PARP1 activation has been demonstrated to be necessary for long-term memory formation, first in *Aplysia* and subsequently validated in mice^48,49^. Additional work using microinfusion of a PARP1 inhibitor into the hippocampus validated these findings and also showed that PARP1 inhibition blocks activity-induced expression of immediate early genes (IEGs) which are necessary for downstream neurotrophic gene expression programs^50^. These independent studies indicate the PARP1 activity is essential for memory formation. An especially important function of PARP1 is how it works in concert with HPF1 to initiate trans-ADP-ribosylation of histones and other non-self proteins which implicates PARP1 as a coordinator of epigenetic chromatin remodeling, DNA damage repair, and transcriptional activation^51–53^. When in complex, the PARP1-HPF1 active site preferentially ADP-ribosylates histone serine residues^54–56^. HPF1 controls the rate of PARP1-mediated histone serine mono-ADP-ribosylation (MARylation)^57^. These serine residues are adjacent to histone lysine residues that are known to be acetylated during memory formation (*i*.*e*. H3K9 and H3K27)^58^, suggesting coordination between histone acetylation and histone ADP-ribosylation in memory formation.

To test this hormetic model of exercise-mediated neuroprotection, we investigated whether acute aerobic exercise induces memory formation concurrent with PARP1 activation and examined the temporal (and spatial) dynamics of exercise-induced gene expression and DNA SSBs primarily in the hippocampus.

## Materials and Methods

### Acute treadmill exercise

All animal studies were performed using male and female C57BL/6J mice obtained from Jackson Laboratories. Animals were housed under standard conditions with ad libitum access to food and water. All experimental procedures were conducted in accordance with the National Institutes of Health guidelines for the care and use of laboratory animals. Protocols were reviewed and approved by the Institutional Animal Care and Use Committee (IACUC) at University of Miami. Animals were habituated to the treadmill for 10 mins at 3 m/min, 5 days/week for 2 weeks with the shock off (ExerGait treadmill, Columbus Instruments). On the day of the acute exercise protocol, animals were brought to the treadmill room 1 hour prior to exercise. Animals assigned to the exercise group were subject to a progressive exercise protocol as detailed in Figure 8B. Sedentary animals were subject to the treadmill for the same time period (40 mins.) at a speed of 1 m/min. After completion of the exercise or sedentary protocol, animals were returned back to their home cage. Animals were exercised six at a time and the treadmill was cleaned with ethanol in between cohorts.

### Sub-threshold novel object location (NOL) or object location memory (OLM)

Three days prior to the OLM test, animals were habituated to the open field (24cm × 24cm). A total of three 10-minute exploration trials was completed over 3 days. Two hours after completion of either the SED or EX protocol, animals were exposed to two identical objects placed in the open field and allowed to explore the objects for 3 minutes and then returned to their home cage. After a 24-hour break, one of the objects was moved to a new location and the animals were then allowed to explore again for 5 minutes. EthoVisionXT animal tracking software was used to measure distance, velocity, cumulative duration of nose-point in object zones, and nose-point object visit frequency. Discrimination index (time exploring novel object location – time exploring familiar object /total time exploring both objects) was used to quantify test performance.

### Tissue Collection

After a week break from exercise and behavior, animals were exercised a final time to study the acute molecular effects of exercise at multiple time points post-exercise. For the 5 minute time-point, animals were moved immediately from the treadmill to sedation with isoflurane. Once the mouse was anesthetized, blood was collected via cardiac perfusion with heparinized PBS (5 U/mL). Liver, spleen, muscle, and adipose tissue was collected and snap frozen in liquid nitrogen. Finally, the brain was removed and microdissected into prefrontal cortex (PFC), hippocampus (HIP), cerebellum (CER), and remaining cortex followed by snap freezing.

### Tissue processing

the mirVana PARIS kit (Ambion) was used for protein and RNA extraction of the hippocampus according to manufacturer’s protocol. Briefly, ½ of the hippocampus (~20 mg) was homogenized in 250 µL of cell disruption buffer supplemented with 1X protease inhibitor (HALT^TM^ protease inhibitor cocktail 100X, ThermoFisher), 1X phosphatase inhibitor (Halt™ Phosphatase Inhibitor Cocktail, ThermoFisher), and RNase inhibitor (SUPERase●In, Invitrogen). Homogenization was performed in the MM 400 (Retsch) with 1 5 mm stainless steel bead (Qiagen) per 250 µL sample at a frequency of 30 Hz for 30 seconds. The sample was then split in half for final protein and RNA isolation.

### Tissue protein extraction

125 µL lysate was further sonicated in the supplemented cell disruption buffer for 30 second intervals on medium frequency for 2 min (BioRupter UCD-200). After a 10-minute incubation on ice, the lysate was centrifuged at max speed for 1.5 min at 4°C and the soluble supernatant was collected and used for ELISA and western blotting. PFC protein was extracted using M-PER (ThermoFisher) supplemented and homogenized identical to lysate in the mirVana PARIS protocol.

### Tissue RNA extraction

Hippocampal RNA was extracted from the remaining 125 µL lysate according to mirVana PARIS protocol using a column-based approach. The protocol was modified to include DNase digestion on final column eluent (TURBO DNA-free, Invitrogen). RNA was quantified using nanodrop and Qubit RNA HS Assay Kit (Thermo Scientific) and used for RT-qPCR and NanoString gene expression assays.

### Western blotting

30 μg of soluble protein lysate with XT Sample Buffer (BioRad) containing β-mercaptoethanol was boiled at 100℃ for 10 minutes. Precision Plus Protein Dual-Color Standards (BioRad) and boiled samples were added to wells of a 26-well 4-12% Criterion^TM^ XT Bis-Tris Protein Gel (BioRad) and run at 150V for 45 min in XT MES buffer. Protein was transferred onto 0.2 μm PVDF membranes using the Trans-Blot Turbo Transfer Pack (BioRad). Blots were incubated, rocking for 1 hour at room temperature with 5% Blotting-Grade Blocker (BioRad) in 1X TBS (BioRad) with 0.1% TWEEN20 (TBST). Following three, 5-minute TBST washes, primary antibody was added. After overnight incubation at 4℃, blots were washed 3 × 5 min. in TBST before secondary antibody in 5% blotting grade blocker for 1 hour at RT. Clarity Western ECL Substrate (BioRad) or SuperSignal West Femto Maximum Sensitivity Substrate (Thermo Scientific) were used to expose blot signal (LI-COR Odyssey DLx).

### Gene Expression

Relative abundance of RNA species was determined using RT-qPCR. 2 μg of RNA from samples were reverse transcribed into cDNA using the qScript cDNA Synthesis Kit (QuantaBio). 100 ng of cDNA was then added to wells in a 384-well format along with TaqMan Fast Advanced Master Mix (Thermo Scientific) and the desired TaqMan probe/primer, with each reaction in technical duplicate. Plates were then loaded into a QuantStudio Real-Time PCR machine and the TaqMan Fast Advanced program was initiated (Applied Biosystems). The ddCt method was used to analyze the output ^101^. Briefly, we calculated the differences between the Ct values for target and reference genes (Rpl37a) as ΔCt and the difference between the resulting ΔCt and that of the vehicle control (calibrator sample) to obtain the ΔΔCt. Results are presented as fold change (RQ = 2^−ΔΔCt^) for mRNA expression relative the vehicle. Any target with Ct values above 32 were not analyzed or repeated with a higher concentration of cDNA. All genes tested by qPCR in these studies were amplified with Taqman primers from Life Technologies/Thermo Fisher Scientific.

### siRNA silencing and in-cell ELISA

On day 1, N2A cells were plated 3 x10^4 cells/well in a 96 well plate in 100 uL growth media. Reverse transfection was performed with RNAiMAX (ThermoFisher) at a final concentration of 10 nmol/L by adding 20 uL of siRNA-RNAiMAX complex in Opti-MEM to each well. Cells were then incubated for 48-96 hours (2-4 days) with the complex before stimulation with 10uM forskolin or DMSO (0.1%) for 10 minutes. Cells were then washed 1X with cold DPBS before fixing with ice cold 100% MeOH at −20C for 10 minutes. After fixation, cells were washed 3X with cold DPBS and blocked in 1X Fish Gel (Rockland Biosciences) diluted in TBS-T (0.1% Tween-20) for 5 hours. Cells were washed 1X in TBS-T before incubation in primary antibody. All primary antibodies were diluted in 1% BSA in TBS-T (antibody buffer) 1:400 and incubated overnight at 4C for 18 hours. The wells were washed 3X with TBS-T and endogenous peroxidase activity was quenched using 3% H2O2 in DPS for 10 minutes. Cells were washed 1X with TBST and then incubated with 1:2000 secondary antibody for 2 hours at room temp in antibody buffer. 100 uL Chromogen was added to each well for 15 minutes followed by 100 uL of stop solution. Absorbance was read at 450 nm using the Envision (Perkin Elmer).

### Statistical Analyses

All data are represented as mean ± SEM and sample size is reported in figure legends. Group means ± SEM and samples sizes (n) are reported in each figure legend. Data were statistically significant if p < 0.05. For all figures, all statistically significant group differences are labeled. For any given group comparison, the lack of any indication of significant difference implies lack of significance by the applied statistical test.

GraphPad Prism software was used for all statistical analyses except those associated with the NanoString assay. Outlier evaluation for all statistical tests was performed using the ROUT method (Q = 1%) in GraphPad Prism

### Tissue preparation and staining

Formalin-fixed paraffin-embedded (FFPE) mouse brain tissues were deparaffinized, rehydrated, and subjected to antigen retrieval procedures. Single-strand DNA breaks were detected using the sSTRIDE assay (Kordon et al., 2020). Briefly, modified nucleotides were enzymatically incorporated at DNA break sites, followed by detection with primary antibodies, secondary antibodies conjugated to oligonucleotides, and rolling circle amplification with fluorescent probe hybridization. Neuronal nuclei were identified with anti-NeuN immunofluorescence, and nuclei were counterstained with DAPI (Abcam ab104139).

### Confocal imaging and analysis

Samples were imaged by confocal microscopy (405 nm, 488 nm, 561 nm excitation). STRIDE foci were quantified using an in-house algorithm that segmented DAPI-stained nuclei, identified foci, and classified neuronal cells based on NeuN positivity. Only foci within NeuN^+^ nuclear masks were analyzed.

### sSTRIDE Quality control and statistics

Segmentation and foci detection were validated by visual inspection, with reanalysis where necessary. Statistical analysis employed generalized linear mixed models (GLMM) to account for nested data (cells within fields of view and samples).

## Results

### Age-related differences in DNA single-strand break distribution reveal distinct patterns

To investigate the physiological distribution of SSBs across genomic elements and their relationship to age, we performed genome-wide SSB mapping in young and aged mouse hippocampus brain. Young mice exhibited significantly higher SSB rates in promoter regions (3.4% vs 2.1%, p=0.0042) and transcription start sites (2.2% vs 1.3%, p=0.0037) compared to aged mice (Figure 1A). Genomic enrichment analysis revealed that SSBs in young mice were significantly enriched at enhancer elements (p=0.006776) and promoter regions (p=0.020371), while showing relative depletion at introns and other non-regulatory sequences (Figure 1B). Gene ontology analysis of SSB-targeted genomic regions in young mice revealed significant enrichment for biological processes including neuronal development, synaptic plasticity regulation, learning and memory pathways, and Wnt signaling, all of which are critical for neuroplasticity and cognitive function (Figure 1C). These findings demonstrate that SSB accumulation follows distinct genomic patterns that change with age, with young animals showing preferential SSB occurrence at transcriptionally active regulatory elements associated with neuroplasticity-related genes.

**Figure 1:**
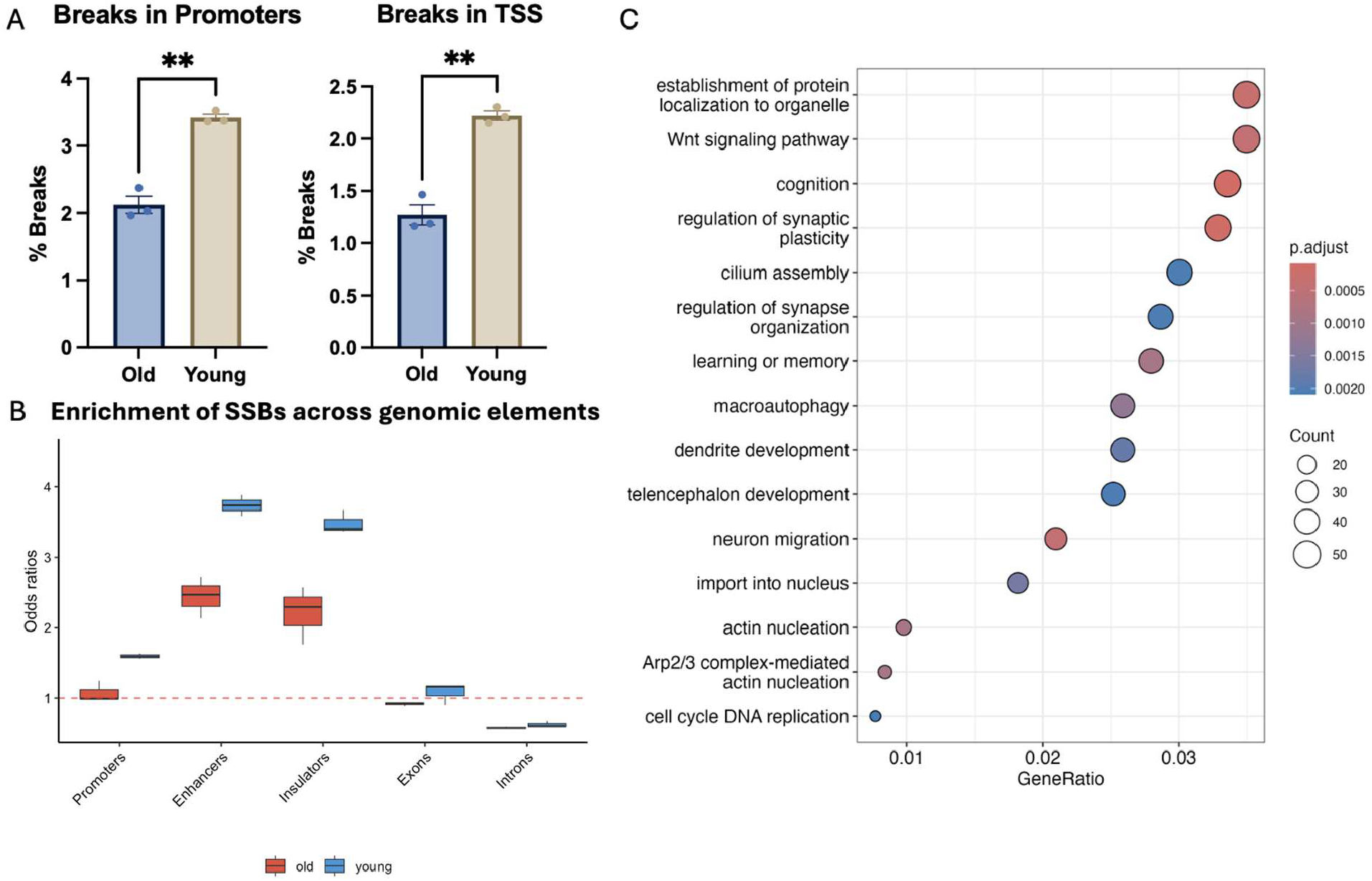
Age-dependent patterns of DNA single-strand breaks reveal preferential targeting of transcriptionally active genomic regions in young mice. **A**. Quantification of SSBs in promoter regions (defined as ±1000 bp from TSS) and transcription start sites (defined as ±500 bp from TSS) shows significantly higher break rates in young compared to old mice. Statistical significance determined by unpaired t-test with Welch’s correction (promoters: p=0.0042; TSS: p=0.0037). **B**. Genomic enrichment analysis demonstrates that SSBs are significantly enriched at enhancers, promoters and insulators in young mice while showing depletion at exons and introns. Enrichment calculated as odds ratios using binomial test for observed vs. expected frequencies based on genomic coverage. The dashed red line indicates odds ratio = 1 (no enrichment). Statistical significance determined by Wilcoxon rank-sum test comparing old vs. young groups (promoters: p=0.020371; enhancers: p=0.006776; insulators: p=0.020935). **C**. Gene ontology analysis of genomic regions targeted by SSBs in young mice reveals significant enrichment for biological processes involved in neuronal development, synaptic plasticity, learning and memory, and Wnt signaling pathways. Circle size indicates gene count; color intensity represents adjusted p-value significance. **p<0.01, *p<0.05. n=3 per age group.

### A single bout of acute aerobic exercise induces sub-threshold memory formation in adult C57Bl6/J mice

In order to investigate the role of exercise-induced neuronal DNA damage in memory formation, an exercise and memory assessment protocol was established based on previously published exercise-induced sub-threshold memory protocols (Fig. 2A, B)^59^. Three consecutive days of habituation to the open field showed that all animals acclimated to the test environment and demonstrated no preference for any of the potential object zones (Fig. 2C,D). Two hours after acute treadmill exercise (EX) or mock exercise (SED), animals were allowed to explore two identical objects in the open field for three minutes. As expected, animals did not show a pre-existing bias for either object (Fig. 2E). After 24 hours, one of the objects was moved to a novel location and mice were evaluated on the cumulative duration of time spent exploring the object in the new location versus the unmoved object. Only exercised mice recognized the novel object location, indicating that acute exercise was able to convert a sub-threshold exposure event into a long-term memory as determined by the discrimination index (DI) (Fig. 2F, *p*< 0.01).

**Figure 2:**
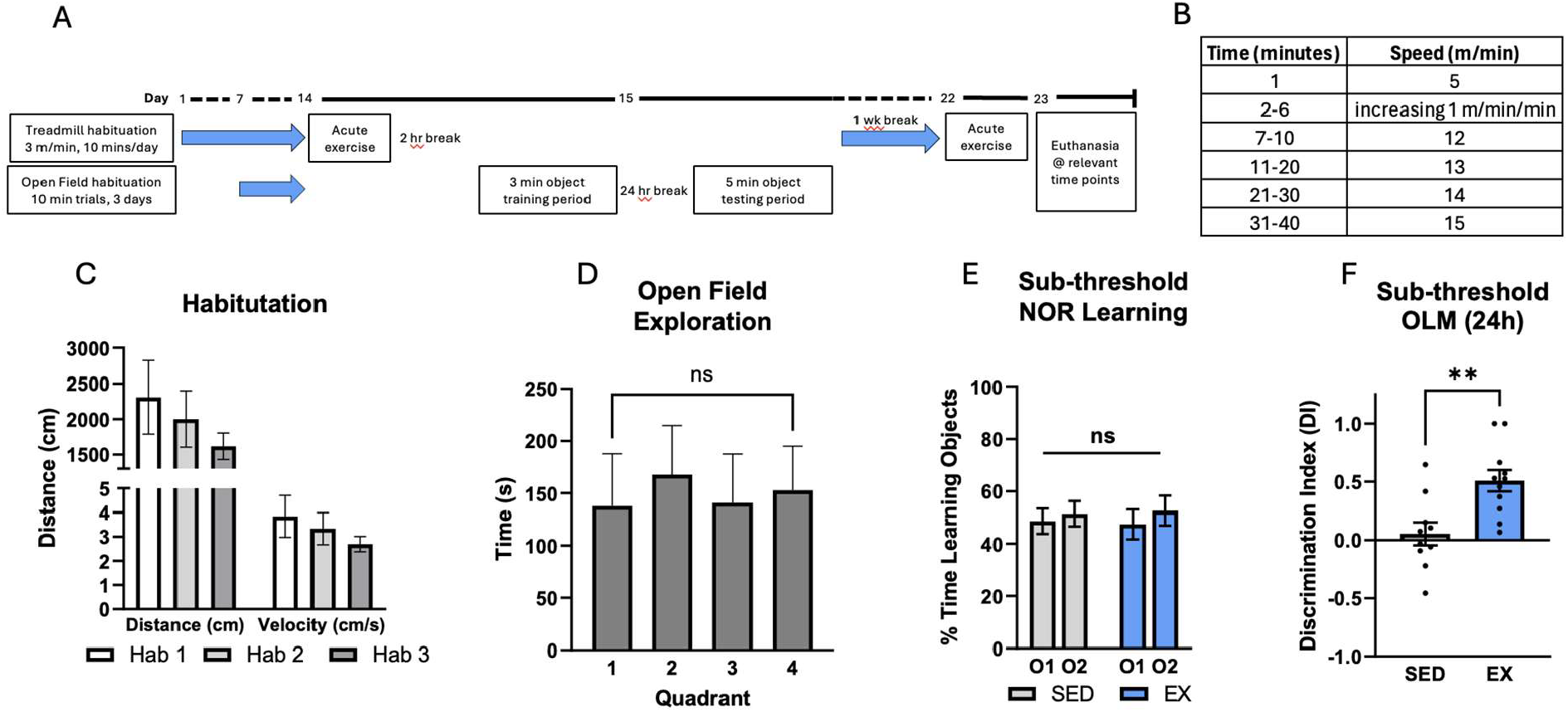
Acute treadmill exercise converts sub-threshold event into long-term memory in adult C57Bl6/J mice. **A**. Experimental timeline of acute exercise habituation, testing, and euthanasia. **B**. Acute exercise treadmill protocol. **C**. Animals habituated to the open field as evidenced by decreased distance and velocity during daily consecutive habituation trials. **D**. Animals did not exhibit a bias for empty object zones during habituation. **E**. Both EX and SED animals explored objects equally during the training period. **F**. EX animals had a significantly higher DI compared to SED animals when exposed to shifted objects 24 hours after initial object training period (n = 11-12/group). Data are represented as mean ± SEM. Two-way ANOVA using Dunnett’s multiple comparisons test or student’s t-test. *p<0.05, **p<0.01.

### Acute treadmill exercise increases neuronal DNA SSBs in the hippocampus

We employed SensiTive Recognition of Individual DNA Ends (STRIDE) technology to spatially visualize post-exercise DNA SSBs across coronal FFPE brain sections in the acutely exercised mice. sSTRIDE is a signal amplification method that directly detects free 3’-OH, indicative of SSBs or intermediates of SSB repair. sSTRIDE is the only available SSB visualization method sensitive and specific enough to directly detect DNA chemistries *in situ*, avoiding reliance on surrogate repair machinery detection and simple immunodetection of DNA chemistries which has well documented limitations in FFPE tissue. Briefly, deoxynucleotide analogues are conjugated to free 3’-OH ends and immunolabeled with 2 unique antibodies. Proximity-restricted rolling circle amplification is initiated between the two unique antibodies, producing a novel DNA sequence that is detected with fluorescent oligomers, revealing the observable signal^60^. In this pilot study, we analyzed sedentary (SED), 5 minute post-exercise mice (5 min.), and 2 hour post-exercise mice (2 hr) for neuron-specific SSBs. Representative images of the sSTRIDe signal in the hippocampus and subsequent quantification show that (1) DNA SSBs accumulate in hippocampal neurons (Fig. 3A), (2) there is a modest increase in sSTRIDE foci per neuronal nuclei immediately post-acute exercise that returns to baseline by 2 hours post-exercise (Fig. 3B), and (3) there is an increase of neuronal nuclei in the 90th percentile (>26 foci) immediately post exercise (Fig. 3C). Based on these data, we continued to investigate neurotrophic and DNA SSB repair-related gene expression *in vivo* and *in vitro*.

**Figure 3:**
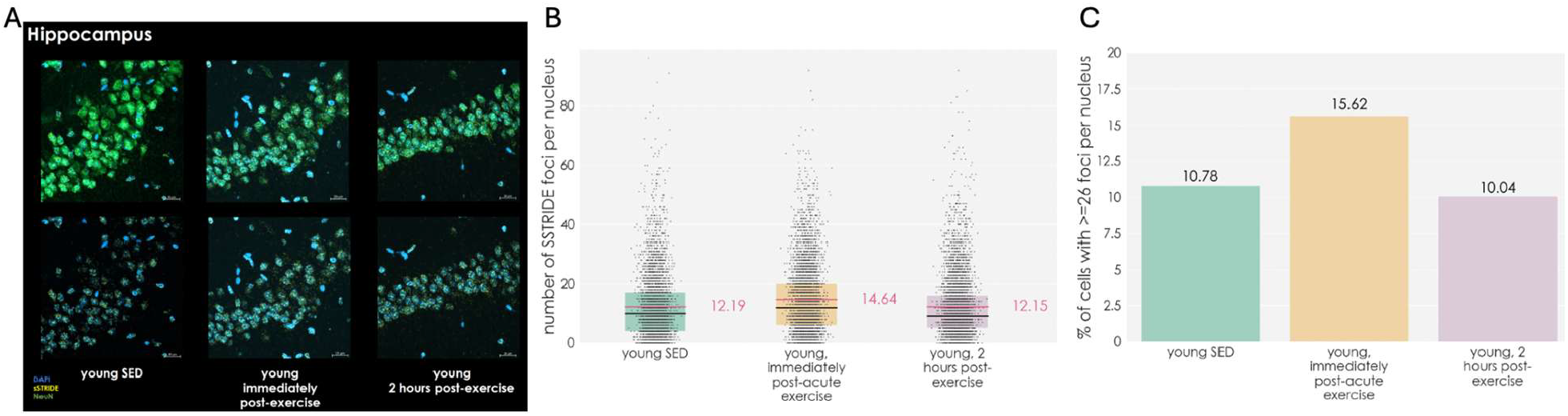
Representative image and quantification of neuronal hippocampal DNA SSBs in acute exercise mice. **A**. Images of hippocampal sections labeled with DAPI (blue), STRIDE (yellow), and NeuN (green) in sedentary, immediately post-acute exercise, and 2 hours post-acute exercise mice. **B**. Quantification of STRIDE foci per nucleus across groups. Each point represents one nucleus. Nuclei from immediately post-exercise mice displayed higher numbers of STRIDE foci (mean = 14.64) compared to sedentary (12.19) and 2 hours post-exercise (12.15). **C**. Proportion of cells with ≥26 STRIDE foci per nucleus, showing a peak immediately post-exercise (15.62%) that returned to sedentary levels by 2 hours post-exercise. Scale bars = 20 μm.

### Acute treadmill exercise upregulates hippocampal neurotrophic gene expression and DNA SSB repair enzymes

After validating that our acute treadmill exercise protocol induces memory formation, we then investigated time-dependent neurotrophic gene expression post-exercise in bulk hippocampal RNA. *Fos*, an activity-dependent immediate early gene (IEG) necessary for *Bdnf* gene expression, was upregulated immediately post-exercise (0 hr: 2.77-fold, *p<*0.001) (Fig. 4A). *Bdnf*, a key mediator of neuroplasticity and neuronal survival, was also found upregulated (2 hr: 1.49-fold, *P<*0.001, 6 hr: 1.41-fold, *p<*0.01) (Fig. 4B). *Creb1*, both an initiating transcription factor and amplifier of *Bdnf* gene expression, was upregulated 1.30-fold post-exercise (*p<*0.05) (Fig. 4C). Lastly, local hippocampal *Igf-1* expression has been shown to support neurogenesis and was upregulated 1.35-fold (*p*<0.05) (Fig. 4D)^61–63^. Immunoblotting of bulk soluble protein from the prefrontal cortex revealed a modest increase in Creb phosphorylation 2 hours post-exercise, further indicating exercise-induced Creb activation in this paradigm (Fig. S1D). Prefrontal cortex Bdnf protein upregulation was not significant (Fig. S1D). These findings corroborate previous reports of exercise-induced neurotrophic signaling and further validate our exercise-induced memory formation behavioral protocol.

**Figure 4:**
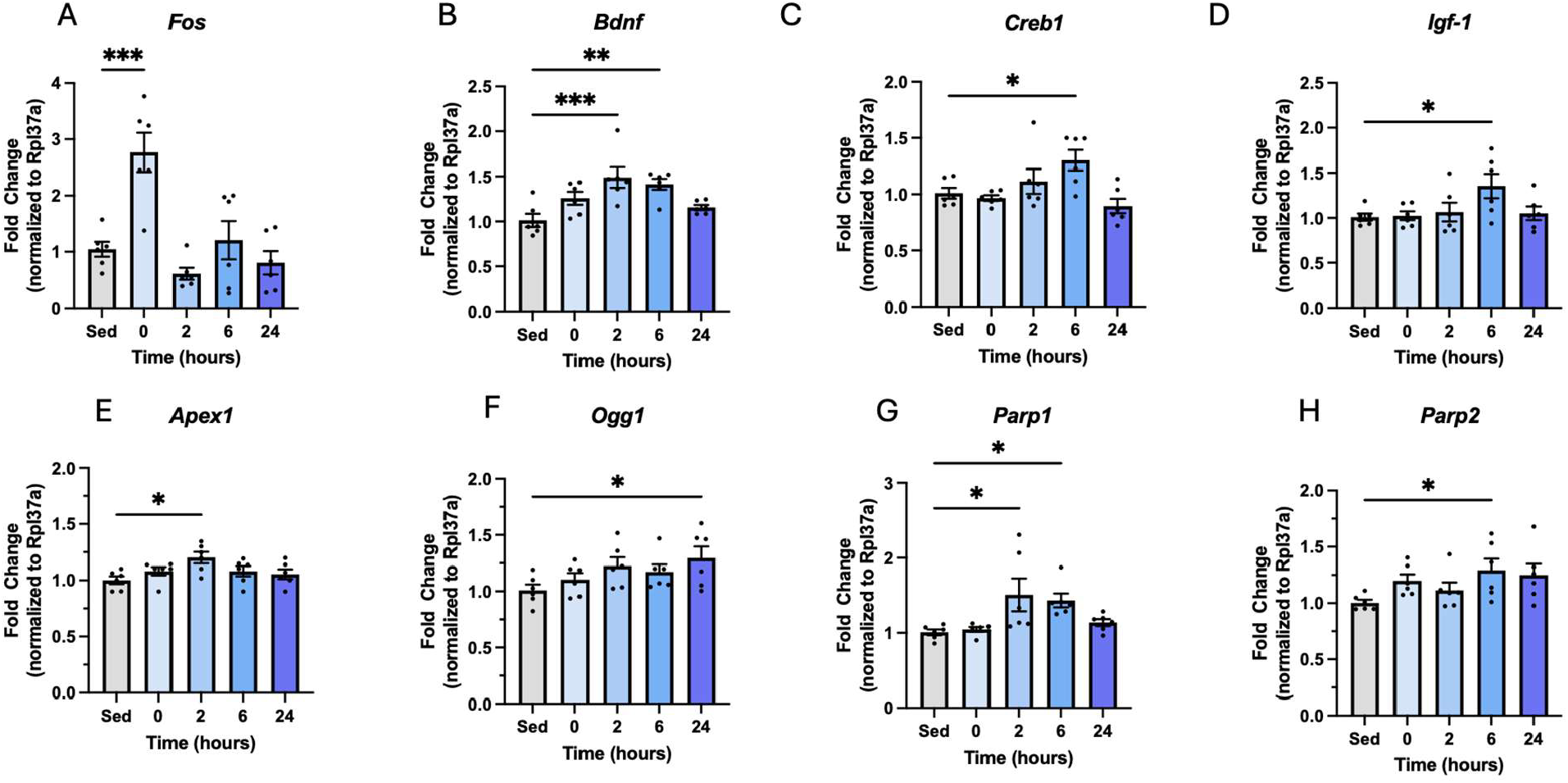
Acute treadmill exercise simultaneously upregulates hippocampal neurotrophic gene expression and DNA SSB repair enzymes. RT-qPCR of bulk hippocampal RNA from the brains of post-exercise mice reveals an upregulation of **A**. *Fos*, **B**. *Bdnf*, **C**. *Creb1*, and **D**. *Igf-1* gene expression at various time-points. In tandem, expression of **E**. *Apex1*, **F**. *Ogg1*, **G**. *Parp1*, and **H**. *Parp2* were also found to be upregulated. (n = 6/timepoint). Data are represented as mean ± SEM. One-way ANOVA using Holm-Sidak’s multiple comparisons test *p<0.05, **p<0.01, ***p<0.001.

We then evaluated the expression of four SSB repair enzymes that have previously been associated with memory to begin to understand the relationship between exercise and DNA damage repair in the brain (Figure 4). *Apex1* (1.21-fold, *p*<0.05) and *Ogg1 (*1.25-fold, *P*<0.05) were found modestly upregulated at 2- and 24-hours post-exercise, respectively (Fig. 4E, F). *Parp1* hippocampal gene expression was upregulated 2- and 6-hours post-exercise (2 hr: 1.51-fold, *P*<0.05, 6-hr: 1.43-fold, *p*<0.05) (Fig. 4G). Additionally, *Parp2* expression was upregulated (1.29-fold, *p*<0.05) at 6 hours post-exercise (Fig. 4H). We did not observe an upregulation of oxidative stress response genes *Cat* or *Sod1* nor hypoxia inducible factor 1 (*Hpf1)* in the HIP in response to acute exercise (Fig. S1A-C). Immunoblotting in the PFC revealed an upregulation of phospho-Creb at 2 hours post-exercise and no change in Bdnf protein expression (Fig. S1D). Plasma H_2_O_2_ and plasma lactate were unchanged post exercise as well as the oxidative stress associated DNA damage marker 8-OH-dG in genomic DNA from the PFC (Fig. S1E-G).

### Hpf1-dependent Parp1 activity is necessary for forskolin- and H_2_O_2_-induced neurotrophic signaling in N2A cells

To further understand the role of Parp1 activity in neurotrophic signaling, siRNA was used to silence Hpf1 in the neuro2a (N2A) mouse neuroblastoma cell model *in vitro*. H_2_O_2_ is a commonly use Parp1 activity inducer as it induces oxidative stress and downstream SSBs. H_2_O_2_ induced Parp1 auto-ADP-ribosylation, Erk2 phosphorylation, and histone 3 (H3) MARylation in N2A cells. These effects were all blocked by 72 hr Hpf1 silencing (Fig. 5A). Using RT-qPCR, we then investigated if Hpf1 silencing blocks *Bdnf* expression induced by forskolin (FSK), a more context-relevant stimulus. In this context, siHpf1 block FSK-induced *Bdnf* expression (*p*<0.05) (Fig. 5B,C). Finally, we used an in-cell ELISA to demonstrate that siHpf1 also blocks 10 minute FSK-induced ERK1/2 phosphorylation but not Creb phosphorylation (Fig. 5D-G). Absorbance signals from the primary antibodies used in the in-cell ELISA were normalized to their respective host secondary only controls (Fig. S2). This is the first time that Hpf1 has been implicated as a mediator of neurotrophic signaling, suggesting that Hpf1-dependent PARP1 activity could be primarily responsible for the role Parp1 plays in memory formation.

**Figure 5:**
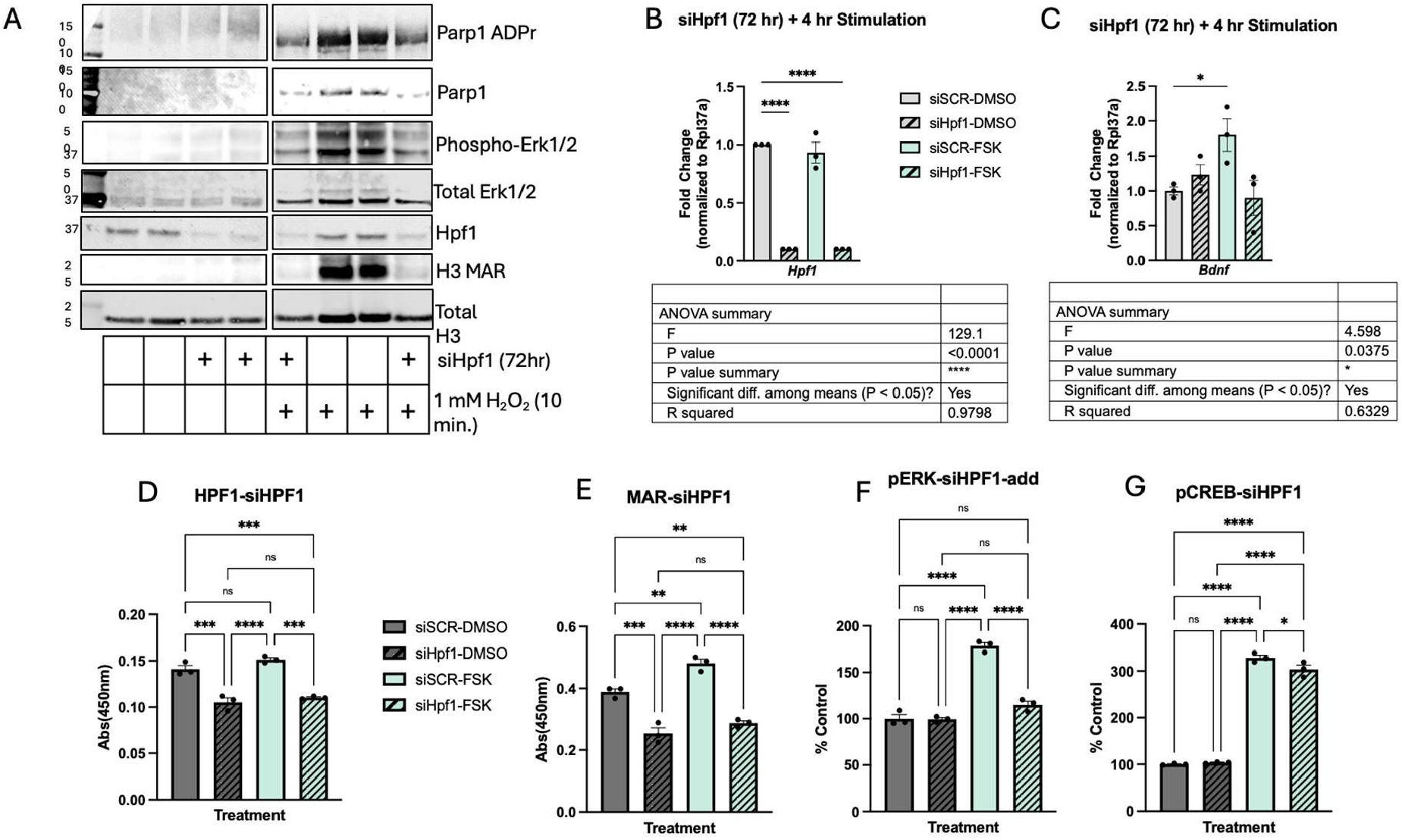
Hpf1-dependent Parp1 activity is necessary for FSK- and H_2_O_2_-induced neurotrophic signaling in N2A cells. **A**. siHpf1 blocks H_2_O_2_-induced Parp1 activation, Erk1/2 phosphorylation, and H3 MARylation. **B**. siHpf1 blocks FSK-induced *Bdnf* gene expression. **C**. siHpf1 blocks FSK-induced MARylation and ERK1/2 phosphorylation but not Creb phosphorylation. (n = 3/group). Data are represented as mean ± SEM. One-way ANOVA using Dunnett’s multiple comparisons test or student’s t-test. *p<0.05, **p<0.01, ***p<0.001, ****p<0.0001.

## Discussion

Our data shows for the first time that acute aerobic exercise upregulates *Parp1* and *Parp2* in the hippocampus of wild type mice in concert with improved long-term memory formation and neurotrophic gene expression. Our *in vitro* mechanistic studies on Parp1 activation implicated Hpf1-dependent Parp1 activation in memory related signaling cascades for the first time. When Parp1 is stimulated with either H_2_O_2_ or FSK, Hpf1 gene expression is unchanged. Studies investigating the effects of increasing the Hpf1:Parp1 ratio show that minor changes in Hpf1 concentration and the Parp1-Hpf1 substrate (NAD+ and ssDNA) concentrations directly influence the switch between Parp1 automodification and trans-ADP-ribosylation of H3^64^. Because of its low stoichiometric relationship with Parp1, its relatively stable expression level in a variety of contexts, and its necessity for initiation of trans-ADP-ribosylation of histone serine residues, Hpf1 is likely a gate keeper of activity-induced chromatin remodeling and an overlooked epigenomic modifier. Chromatin immunoprecipitation (ChIP-seq) studies of Hpf1 have not yet been performed. The ability of Hpf1 to control Parp1 activity has been described as a “hit-and-run” mechanism where Hpf1 quickly associates and disassociates from an individual Parp1-Hpf1-nuclesome-NAD+ complex with a half-life of about 10s^57^. Because of this quick on-off interaction, time-resolved capture of the stimulated ADP-ribosylome in neurons may reveal more useful information for therapeutic discovery. The ADP-ribosylome has so far only been mapped in HeLa cells^65^. ADP-ribosylhydrolase (ARH3) is the only known enzyme that directly reverses serine MARylation. Defects in this enzyme lead to a genetic disease known as childhood-onset, stress-induced, with variable ataxia and seizures (CONDSIAS)^66^. Defective ARH3 leaves ADP-ribose “scars” on H1 and H3 that inhibit the ability of the chromatin to return to its un-stimulated ground state. This finding directly aligns with our hypothesis that the Parp1-Hpf1 complex is a direct mediator of exercise (stress)-induced neuronal signaling. Histone ADP-ribose “scarring” could be a novel and more representative pathological hallmark of aging and neurodegenerative disease since the deposition and removal of MAR is tightly regulated by two, well-defined opposing enzyme complexes. This is as opposed to amyloid and tau pathology (slow and unknown etiology) or DNA methylation (multifactorial regulation). The extent to which ADP-ribosylation in the periphery reflects the state of the CNS is also completely unknown.

## Limitations

A few challenges listed below limit the definitive conclusions about exercise-induced DNA damage and memory formation. However, the observations from the current study strongly suggest a hormesis-type role of SSB DNA damage for a healthy brain (young vs. aged) and highlight the benefits of exercise for DNA damage repair and memory formation. More work targeting the identified pathways is needed. Translation Challenges: Exercise is inherently a multi-organ intervention that cannot be fully replicated in vitro, requiring researchers to use isolated tool compounds that may not capture the complete physiological response. Additionally, mouse models have substantially different metabolisms and DNA repair kinetics compared to humans, particularly regarding aging-related diseases, limiting translational relevance. However, mice allow experiments in a more controlled environment, guiding future work in humans. Integration with human data is needed. In Vitro Model Limitations: The study used artificial PARP1 activators (forskolin, H2O2) that only partially mimic exercise-induced signaling. Forskolin directly activates signaling pathways, obscuring the natural relationship between HPF1 and memory formation. The N2A cell model lacked appropriate serotonin receptors, preventing use of more physiologically relevant stimulants. Temporal Complexity: Exercise effects on the brain are subtle and accumulate over time, using the SSiNGLe methods and sSTRIDE to capture both acute and chronic changes in different age groups would be beneficial. These limitations underscore the need for future studies with direct SSB mapping, improved spatial transcriptomics, and better in vitro models to validate the hormetic DNA damage hypothesis.

## Future directions

As previously mentioned, the priority of future experiments will be to establish the exercise induced SSB signature of gDNA in the PBMCs and hippocampal neurons of acutely exercised mice at the previously established post-exercise timepoints. Additionally, this DNA will also be used to perform bisulphite sequencing to contextualize the SSBreakome with the more established DNA methylome which has previously been linked to aging and health status^228^. Another major goal of this work is to link the SSBreakome with the exercise-induced transcriptional response. This will allow us to unravel the functional consequences of exercise-induced SSBs. RNA sequencing in PBMCs and a repeat of spatial transcriptomics in the brains of acutely exercised animals will be essential to accomplish this.

Beyond establishing a connection between DNA SSBs and gene expression, probing the role of Hpf1-dependent Parp1 activity in neurotrophic signaling *in vitro* and *in vivo* will reveal if histone ADP-ribosylation is an essential mechanism for memory formation. This will be done by silencing Hpf1 *in vitro* and creating a dominant-negative Hpf1 cell line and investigating the effects of Parp1 stimulants on downstream neurotrophic signaling. Finally, the same acute exercise experiments will be performed with an Hpf1 knockout mouse model that has already been generated in the laboratory of Dr. Ivan Ahel but not yet phenotyped in this context to understand the role of Hpf1 in memory formation in a mammalian model.

## Acknowledgments

We would like to thank the Florida Department of Health Ed and Ethel Moore Alzheimer’s Disease Research Program for support through grant # 22A14 awarded to CW and CHV.

## Notes

### Competing Interest Statement

The authors have declared no competing interest.

