## Supplementary figures for "Exercise-induced DNA damage response and memory formation in mice"

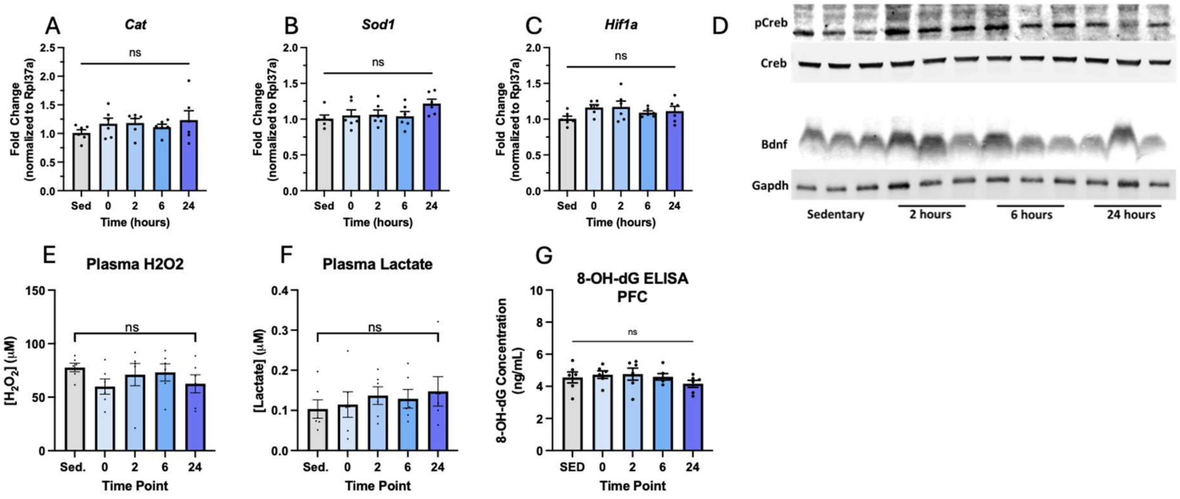


**Supplementary Figure 1. A-C.** mRNA expression levels of Cat, Sod1, and Mfn2 in tissue samples measured by qPCR at 0 (sedentary), 3, 7, and 24 hours post-exercise. Data are normalized to housekeeping genes and expressed as fold change relative to sedentary controls. **D.** Representative Western blot images showing levels of phosphorylated and total Crb, Mfn2, and loading control across the indicated time points. **E.** Plasma levels of hydrogen peroxide ($H_2O_2$). **F.** Plasma lactate concentrations. **G.** Quantification of 8-OHdG levels in PFC tissue measured by ELISA. Data are presented as mean ± SEM. Statistical significance is indicated by ns = not significant.

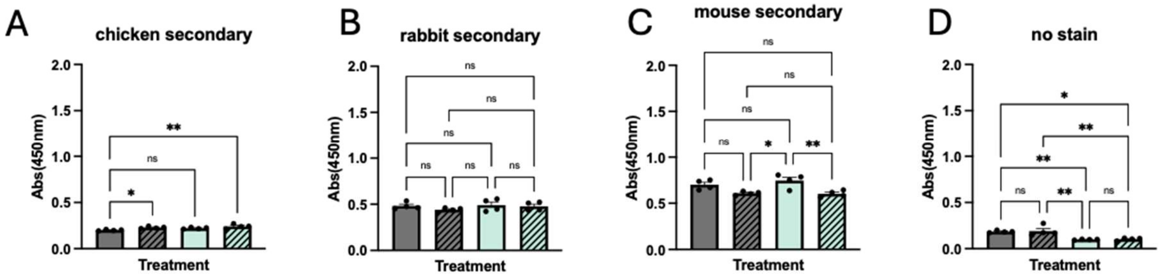


**Supplementary Figure 2. A-D.** Absorbance (450 nm) measurements from in-cell ELISA assays showing background signal across different treatment conditions using chicken (A), rabbit (B), and mouse(C) host species secondary antibodies alone, as well as a no antibody control (D). Absorbance readings were normalized to each respective secondary-only control condition. Data are presented as mean ± SEM. Statistical significance was determined by one-way ANOVA with multiple comparisons: $p < 0.05$, $p < 0.01$, ns = not significant.
